# Information Bottleneck Dominates Adversarial Training for Ancestry-Invariant Polygenic Risk Prediction: Dimensionality, Not Gradient Reversal, Controls the Fairness-Accuracy Tradeoff

**DOI:** 10.64898/2026.04.24.720752

**Authors:** Philip Phuong Tran, Anh T. Do

**Affiliations:** Paragon Biosignals (Univault Technologies LLC), Salt Lake City, Utah, USA

**Keywords:** information bottleneck, adversarial fairness, polygenic risk scores, ancestry bias, representation learning, gradient reversal, latent dimensionality, cross-ancestry prediction

## Abstract

In adversarial representation learning for fair prediction, the gradient reversal coefficient (*λ*) is widely treated as the primary control for sensitive-attribute invariance. We show this assumption is wrong. Using a dual-stream architecture for cross-ancestry polygenic risk score (PRS) prediction, we demonstrate that **latent dimensionality** — the information bottleneck — accounts for 8–27 × more variance in ancestry leakage than adversarial strength. Varying *λ* across a 20 × range changes leakage by only 2.2 percentage points; varying dimensionality across a 16 × range changes it by 46.6 pp. At dimension 8 with *no adversarial training* (*λ* = 0), ancestry leakage is 32.9% (chance = 20%): the bottleneck alone achieves near-invariance. The adversary architecture (linear vs deep MLP) is equally irrelevant (0.6 pp range). We validate this finding across two unrelated domains — genomic ancestry invariance (6 clinical traits, 1000 Genomes, *n* = 2,504) and EEG subject invariance (pretrained HFTP + Braindecode dual-domain model, 20 subjects) — observing consistent dimensionality dominance (12.7:1 ratio in EEG).

For the genomic application, Stream 1 encodes population structure via DCT-II frequency-domain features (136 coefficients); Stream 2 encodes phenotype signal from top PRS SNPs (PCA to 128 dimensions). The architecture works equally well with standard genomic PCA as the ancestry stream (*R*^2^ = 0.217 vs 0.222), confirming the contribution is architectural, not encoding-specific. African-ancestry PRS reconstruction *R*^2^ improves on all six traits (e.g., +5.1 pp for coronary artery disease). Linear models achieve higher aggregate *R*^2^ but fail catastrophically on cross-ancestry transfer (*R*^2^ = − 12.45 for African-ancestry CAD). We emphasize that we predict PRS (a computed score), not disease phenotypes; validation on biobank-scale phenotype data is ongoing.

These results suggest the adversarial fairness community has been over-investing in adversary engineering relative to simple capacity control. Practitioners should select latent dimensionality first to set the information budget for the fairness-accuracy tradeoff, then optionally use adversarial training for marginal refinement.

## 1 Introduction

### 1.1 The PRS Equity Problem

Polygenic risk scores aggregate the effects of thousands to millions of genetic variants into a single quantitative measure of disease risk, and have emerged as a cornerstone of genomic medicine for risk stratification of common diseases [Khera et al., 2018]. However, PRS exhibit a well-documented and clinically consequential failure: they perform substantially worse when applied to populations whose ancestry differs from the discovery cohort. Martin et al. [2019] demonstrated that PRS accuracy degrades by up to 78% when European-derived scores are applied to African-ancestry populations, with intermediate degradation for other non-European ancestries. This disparity is not academic — it means that the genomic tools being integrated into clinical decision-making systematically fail for the populations that already face the greatest health disparities.

The root cause is well understood: GWAS effect sizes are estimated in populations with specific allele frequency distributions and linkage disequilibrium (LD) patterns. When these estimates are applied to populations with different allele frequencies and LD structure, the PRS captures a mixture of disease-relevant signal and ancestry-correlated noise [Ding et al., 2023]. Standard PRS computation cannot separate these components, though ancestry deconvolution approaches have shown partial improvements in admixed individuals [Marnetto et al., 2020].

### 1.2 Current Approaches and Their Limitations

Several approaches have been proposed to address PRS transferability. Multi-ancestry GWAS meta-analyses increase representation but require large sample sizes from diverse populations that remain scarce [Ruan et al., 2022]. Statistical methods such as LDpred2 [Privé et al., 2021] and PRS-CSx [Ruan et al., 2022] model LD differences across populations but operate on summary statistics rather than individual-level representations. Transfer learning approaches adapt European-trained models to target populations but still require ancestry-matched calibration data.

These methods share a common strategy: they attempt to correct ancestry bias through statistical adjustment of weights or calibration. An alternative strategy, which we explore here, is to address ancestry confounding at the *representation level* — learning features that explicitly separate ancestry information from phenotype-relevant signal before prediction.

### 1.3 Key Idea: Architectural Disentanglement

We propose a dual-stream architecture that separates ancestry and phenotype information into distinct latent representations using adversarial training. Rather than attempting to remove ancestry from a single feature set, we provide the model with two input streams:

- **Stream 1 (Ancestry)**: Frequency-domain features computed via DCT-II encoding of genotype data across 22 chromosomes. These features capture population structure (82% ancestry classification accuracy from 136 coefficients) and serve as a strong ancestry signal for the adversary.
- **Stream 2 (Phenotype)**: Dosage features for the top-weighted PRS SNPs, reduced via PCA. These features carry the phenotype-relevant signal that standard PRS aims to capture.

A gradient reversal layer (GRL) [Ganin et al., 2016] is applied specifically to Stream 2 before it reaches the adversary, forcing the phenotype encoder to discard ancestry information. Critically, Stream 1 passes to the adversary without reversal, providing an explicit ancestry reference that makes the adversary stronger — and therefore the disentanglement more effective — than single-stream approaches where the adversary must discover ancestry signal from the same features it is trying to remove it from.

The architectural principle of conditioning a domain adversary on a privileged sensitive-attribute encoding has precedent in the fair representation learning literature [Creager et al., 2019]. Our specific contribution is the application of this principle to genomic PRS prediction, with the DCT-II encoding providing the privileged ancestry reference, and the empirical demonstration that this architecture generalizes across multiple disease traits. The design was motivated by prior work applying dual-domain architectures with gradient reversal to subject-invariant biosignal classification [Tran and Do, 2026].

### 1.4 Contributions

1. The empirical finding that **latent dimensionality dominates adversarial training strength** (8–27 ×) for controlling sensitive-attribute invariance in entangled-feature settings, with adversary architecture contributing negligibly (0.6 pp range across four architectures). This directly contradicts the prevailing practice of tuning *λ* as the primary fairness control.
2. **Cross-domain validation** of this finding in two unrelated domains — genomic ancestry invariance (6 traits) and EEG subject invariance (pretrained dual-domain model) — demonstrating consistent dimensionality dominance with a 12.7:1 ratio in EEG.
3. A dual-stream adversarial architecture for PRS prediction that explicitly disentangles ancestry and phenotype representations, with a dedicated ancestry stream that strengthens the adversary. The architecture is encoding-agnostic: standard genomic PCA and DCT-II produce equivalent results.
4. Demonstration that linear models fail catastrophically on cross-ancestry transfer (*R*^2^ = − 12.45) while the dual-stream architecture degrades gracefully, with implications for deployment robustness under ancestry distribution shift.

We emphasize that this is a proof-of-concept study on *n* = 2,504 individuals from 1000 Genomes. We validate the *method*, not its clinical PRS performance at biobank scale. Large-scale validation is ongoing.

## 2 Methods

### 2.1 Dataset

We use the 1000 Genomes Project Phase 3 dataset [1000 Genomes Project Consortium, 2015], comprising 2,504 individuals from 26 populations grouped into 5 super-populations: African (AFR; *n* = 661), Admixed American (AMR; *n* = 347), East Asian (EAS; *n* = 504), European (EUR; *n* = 503), and South Asian (SAS; *n* = 489). Whole-genome variant call data for all 22 autosomes were obtained in VCF format and converted to plink2 binary format (pgen/pvar/psam) for efficient genotype access.

### 2.2 PRS Targets

PRS weights for six traits were obtained from the PGS Catalog [Lambert et al., 2021] (Table 1). For each trait, per-individual PRS was computed as the weighted sum of effect allele dosages using published effect weights. These PRS values serve as prediction targets; we do not have individual-level phenotypes in 1000 Genomes. We note that PRS targets are themselves ancestry-biased — our method addresses ancestry bias in the *model* but cannot correct bias in the *weights*.

**Table 1:** Traits and PRS weights used in this study. All weights are from the PGS Catalog, harmonized to GRCh37.

| Trait | PGS ID | Variants | Source |
| --- | --- | --- | --- |
| Height | PGS002146 | ~2M | Yengo et al. 2022 |
| Type 2 Diabetes (T2D) | PGS000014 | 6,917,436 | Khera et al. Nat Genet 2018 |
| Coronary Artery Disease (CAD) | PGS000018 | 1,745,180 | Khera et al. 2018 |
| Body Mass Index (BMI) | PGS000027 | 2,100,302 | Khera et al. 2018 |
| Atrial Fibrillation (AF) | PGS000016 | 6,730,541 | Khera et al. 2018 |
| Breast Cancer (BrCa) | PGS000004 | 313 | Mavaddat et al. AJHG 2018 |

### 2.3 Feature Construction

#### 2.3.1 Stream 1: Ancestry Features (DCT-II Encoding)

Following the General Learning Encoder (GLE) framework [Tran and Do, 2026], we encode genotype data into frequency-domain representations using the Discrete Cosine Transform Type II. For each chromosome *c* ∈ {1, …, 22}, the variant dosage vector for each individual is transformed via DCT-II, and the mean across chromosomes yields a 136-dimensional feature vector per individual:

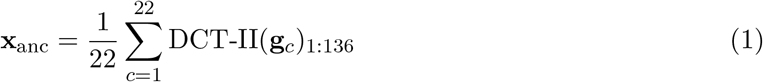

where **g**_*c*_ is the genotype dosage vector for chromosome *c* and the subscript 1:136 denotes retention of the first 136 coefficients. This encoding captures allele frequency patterns across the genome in a compact representation: a logistic regression classifier achieves 81.2% ancestry classification accuracy from these 136 features alone (chance = 20%). The DCT-II is deterministic and does not require fitting to data, unlike PCA [Price et al., 2006].

We note that the DCT-II transform applied to genotype dosages is unconventional in genomics. Standard genomic PCA [Price et al., 2006] achieves substantially higher ancestry classification accuracy (96.6% vs 81.2% for DCT-II at the same dimensionality). We do not claim that DCT-II is superior to PCA for ancestry encoding. Our motivation for DCT-II is practical: as a deterministic, fixed transform, it avoids data leakage that can occur with PCA computed on training data and applied to test data, and it enables privacy-preserving architectures where the encoding can be computed on-device without sharing raw genotype data. A comparison of dual-stream performance using PCA vs DCT-II ancestry features is presented in Section 3.5.

#### 2.3.2 Stream 2: Phenotype Features (PRS SNP Dosages)

For each trait, we extract genotype dosages for the 2,000 SNPs with the largest absolute effect weights in the corresponding PGS weight file. Dosages are read directly from pgen files using pgenlib, with allele orientation matched to the PGS effect allele. The resulting 2,504 × 2,000 dosage matrix is reduced to 128 dimensions via PCA (fit on training data only within each cross-validation fold to prevent leakage).

PRS for each individual is simultaneously computed as the weighted sum of all matched SNP dosages across 22 chromosomes, serving as the prediction target.

### 2.4 Model Architecture

#### 2.4.1 Dual-Stream GRL

The dual-stream model consists of three components (Fig. 1):

**Figure 1:**
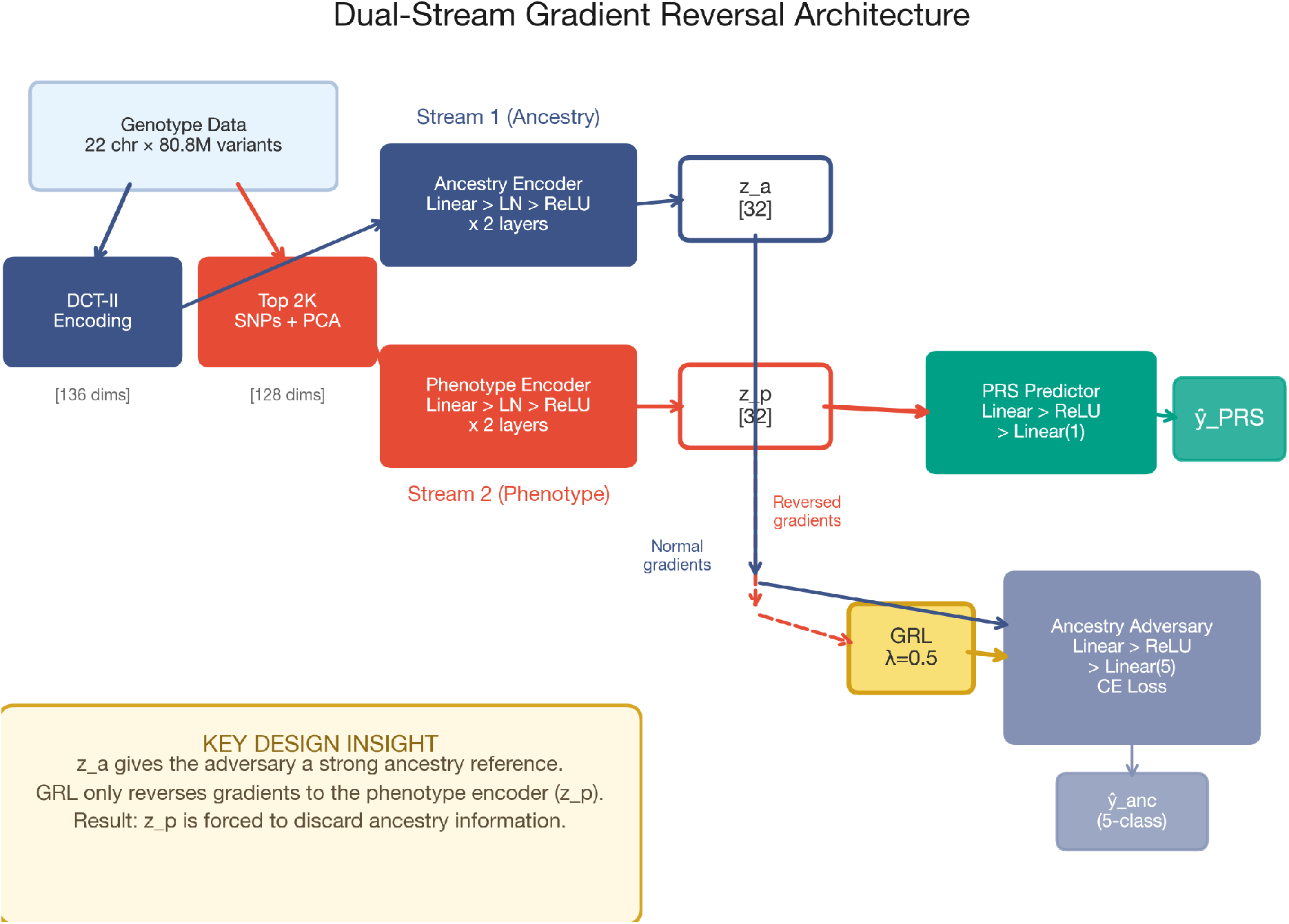
Dual-stream gradient reversal architecture. Stream 1 (blue) encodes population structure via DCT-II features; Stream 2 (red) encodes phenotype-relevant PRS SNP features. The ancestry latent **z**_*a*_ passes to the adversary with normal gradients, providing a strong ancestry reference. The phenotype latent **z**_*p*_ passes through a gradient reversal layer (GRL), forcing the phenotype encoder to discard ancestry information. The predictor operates only on **z**_*p*_.

**Figure 2:**
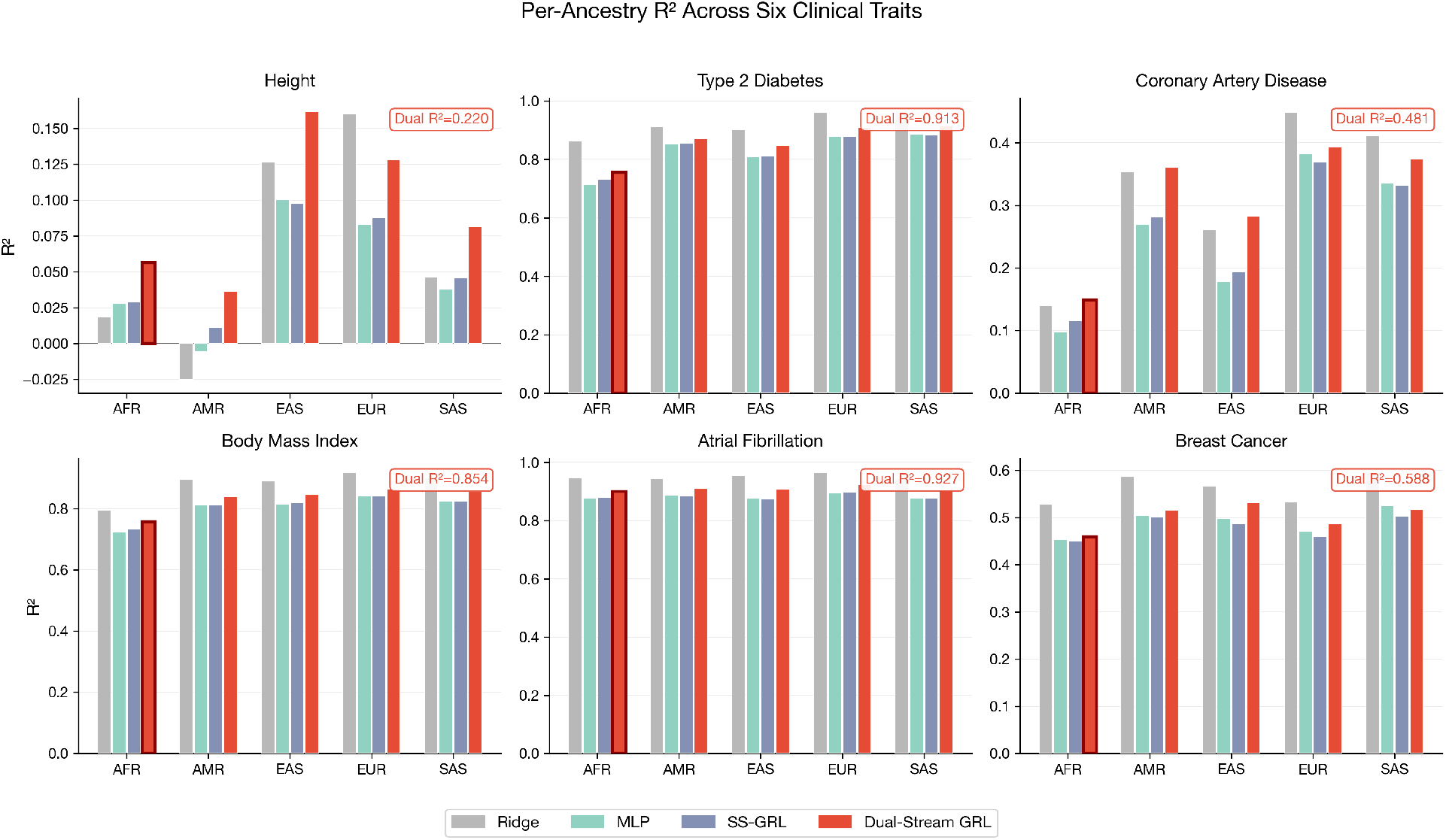
Per-ancestry *R*^2^ across six clinical traits for all four models. The dual-stream GRL (red) consistently outperforms single-stream neural baselines (MLP, SS-GRL) across ancestries. AFR bars for the dual-stream model are highlighted with dark borders. Ridge (gray) achieves the highest per-ancestry *R*^2^ on strong-signal traits (T2D, AF) but fails on cross-ancestry transfer (Fig. 3).

**Figure 3:**
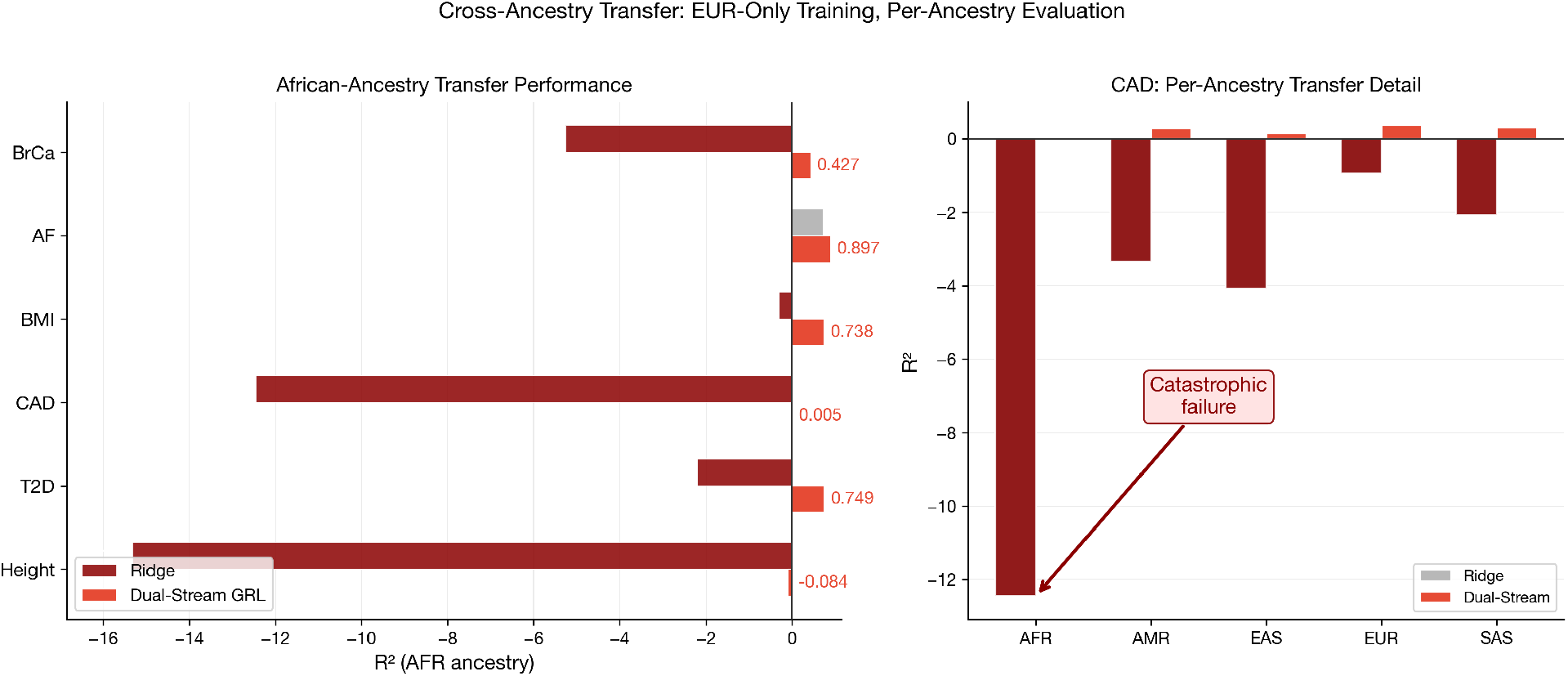
Cross-ancestry transfer robustness. **Left:** African-ancestry *R*^2^ when models are trained on EUR-only data. Ridge regression (dark red) produces deeply negative *R*^2^ on 5 of 6 traits, indicating predictions worse than the population mean. The dual-stream GRL (red) maintains positive *R*^2^ on 5 of 6 traits. **Right:** Per-ancestry detail for CAD (worst case), showing Ridge collapse across all non-EUR ancestries.

**Figure 4:**
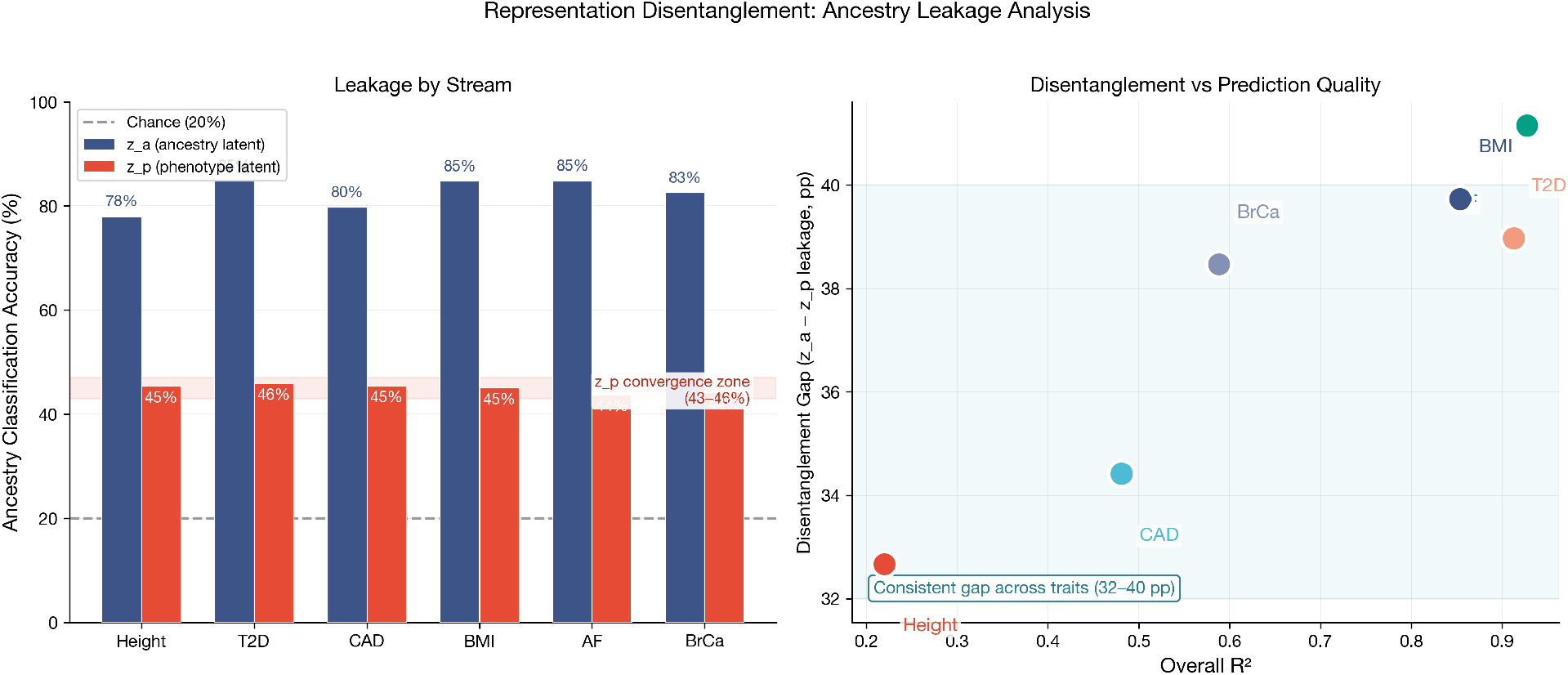
Representation disentanglement analysis. **Left:** Ancestry classification accuracy from **z**_*a*_ (blue, 78–85%) vs **z**_*p*_ (red, 43–46%) across all six traits, with chance level at 20% (dashed). The **z**_*p*_ convergence zone (shaded) shows that phenotype latent leakage is remarkably consistent regardless of trait. **Right:** Disentanglement gap (**z**_*a*_ − **z**_*p*_ leakage) vs overall *R*^2^. The gap remains in a narrow 32–40 pp band across traits that vary widely in prediction difficulty.

**Figure 5:**
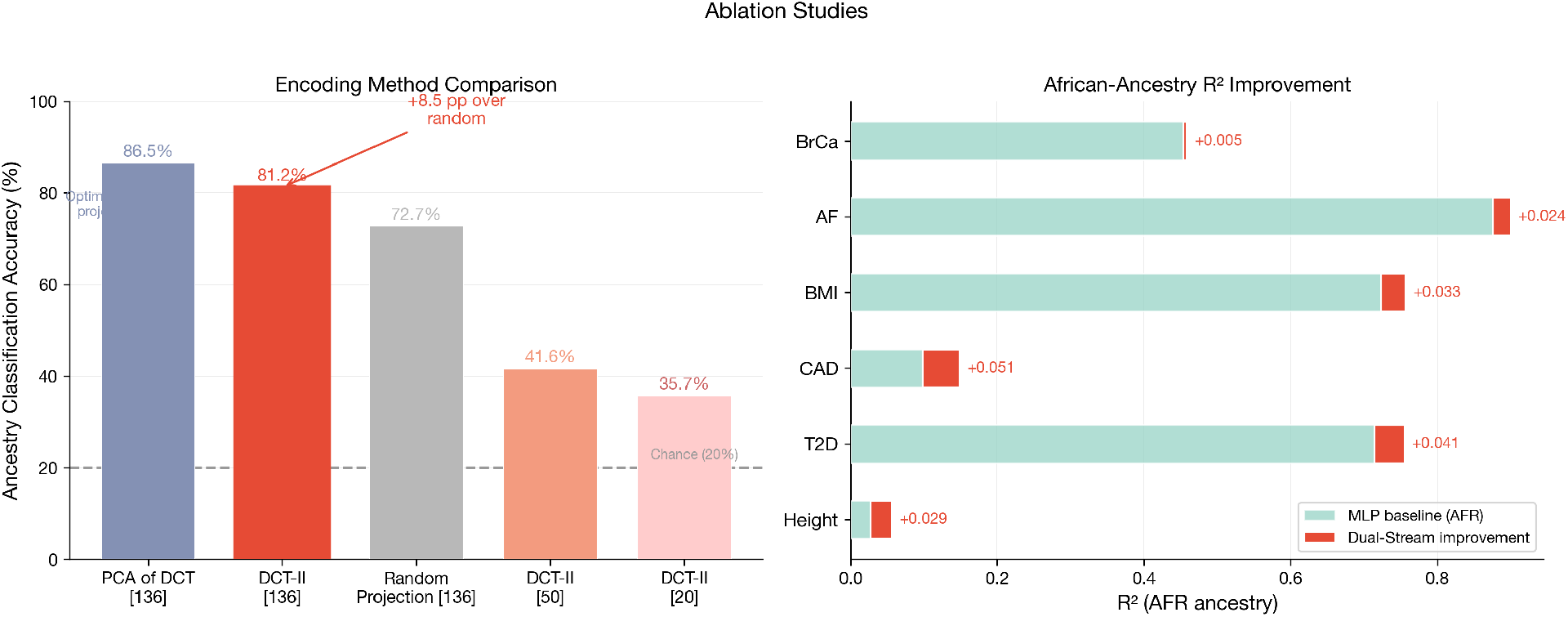
Ablation studies. **Left:** Ancestry classification accuracy from different genotype encodings. Genomic PCA (not shown, 96.6–98.6%) substantially outperforms DCT-II (81.2%), which in turn outperforms random projection (72.7%). **Right:** African-ancestry *R*^2^ improvement of the dual-stream GRL (red) over MLP baseline (green) across all six traits, showing consistent gains.

##### Ancestry encoder (Stream 1)

**x**_anc_ ∈ ℝ^136^ → Linear(136, 64) → LayerNorm → ReLU → Dropout(0.3) → Linear(64, 32) → LayerNorm → ReLU → **z**_*a*_ ∈ ℝ^32^.

##### Phenotype encoder (Stream 2)

**x**_phe_ ∈ ℝ^128^ → Linear(128, 64) → LayerNorm → ReLU → Dropout(0.3) → Linear(64, 32) → LayerNorm → ReLU → **z**_*p*_ ∈ ℝ^32^.

##### Predictor

**z**_*p*_ → Linear(32, 16) → ReLU → Dropout(0.3) → Linear(16, 1) → ŷ_PRS_. The predictor operates *only* on the phenotype latent **z**_*p*_.

##### Adversary

[**z**_*a*_; GRL(**z**_*p*_)] → Linear(64, 32) → ReLU → Dropout(0.3) → Linear(32, 5) → ŷ_anc_.

The adversary receives **z**_*a*_ with normal gradient flow and **z**_*p*_ through a gradient reversal layer. This is the key design choice: **z**_*a*_ provides the adversary with a strong, direct ancestry signal, making it harder for **z**_*p*_ to “hide” residual ancestry information. The GRL reversal coefficient *λ* is warmed up linearly over the first 20 epochs from 0 to 0.5.

The total loss is:

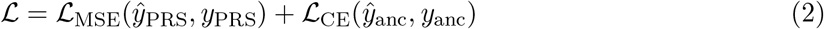

where ℒ_CE_ uses inverse-frequency class weights to handle ancestry imbalance. Due to gradient reversal, the phenotype encoder receives gradients that *oppose* ancestry classification, forcing **z**_*p*_ to be ancestry-uninformative.

The total parameter count is 24,358 — deliberately small to avoid overfitting on *n* = 2,504.

#### 2.4.2 PRS Residualization

To remove trivial ancestry mean differences from the prediction target, we residualize PRS by subtracting per-ancestry means computed on training data only. Predictions are un-residualized before evaluation by adding back the training-set ancestry means. This ensures the model must learn *within-ancestry* PRS variation rather than simply predicting ancestry means.

### 2.5 Baselines

We compare against three baselines, all using the same concatenated feature set (**x**_anc_⊕**x**_phe_ ∈ ℝ^264^):

1. **Ridge regression**: Linear model with *L*_2_ regularization (*α* = 1.0). 264 parameters. Represents the statistical genetics standard.
2. **Single-stream MLP**: Same architecture as one stream of the dual model, applied to concatenated features. 19,777 parameters. Tests whether a neural network alone (without adversarial training) improves over Ridge.
3. **Single-stream GRL**: Single encoder with GRL and ancestry adversary applied to the shared latent. 20,390 parameters. Tests whether dual-stream separation provides benefit over standard domain adversarial training [Ganin et al., 2016].

All neural models use AdamW (lr = 10^−3^, weight decay = 0.01), cosine annealing, gradient clipping (max norm 1.0), and early stopping (patience 25 epochs, max 300 epochs). Training uses Apple MPS acceleration where available.

### 2.6 Evaluation Protocol

We use 5-fold stratified cross-validation (stratified by ancestry) repeated across 5 random seeds, yielding 25 evaluation runs per trait per model. Within each fold, 20% of the training set is held out for early stopping.

**Metrics:**

- *R*^2^ (overall and per-ancestry): coefficient of determination on un-residualized predictions.
- **Ancestry leakage**: accuracy of a logistic regression classifier trained to predict ancestry from latent representations (**z**_*a*_, **z**_*p*_, or **z** for single-stream models). Chance level is 20%.
- **Cross-ancestry transfer**: Ridge regression trained on EUR-ancestry latent features only, evaluated per ancestry. Tests whether the representation generalizes without ancestry-matched training data.

Statistical significance is assessed via paired *t*-tests (or Wilcoxon signed-rank when normality is rejected by Shapiro-Wilk) across the 25 paired runs. Effect sizes are reported as Cohen’s *d*. Bootstrap 95% confidence intervals are computed from 10,000 resamples.

## 3 Results

### 3.1 Multi-Trait Performance

Table 2 presents the primary results across all six traits. The dual-stream GRL significantly outperforms both single-stream neural baselines (MLP, SS-GRL) on five of six traits at *p* < 0.001. BrCa reaches *p* = 0.030, which does not survive Bonferroni correction for six traits (threshold = 0.0083); we retain it in the main analysis as a constrained edge case (only 313 PRS SNPs) but note that this result should be interpreted cautiously.

**Table 2:** Dual-stream GRL performance across six traits. *R*^2^ values are means over 25 runs (5 seeds × 5 folds) with bootstrap 95% CIs (10,000 resamples). Ancestry leakage is classification accuracy from logistic regression on the latent space (chance = 20%). Δ vs MLP shows the paired improvement over the single-stream MLP baseline. BrCa does not survive Bonferroni correction (*p* > 0.0083).

| Trait | Per-Ancestry $R^2$ | | | | | | Leakage | | vs MLP | |
| --- | --- | --- | --- | --- | --- | --- | --- | --- | --- | --- |
| | Overall [95% CI] | AFR [95% CI] | AMR | EAS | EUR | SAS | $z_a$ | $z_p$ | $\Delta R^2$ | $p$ |
| Height | .220 [.203,.235] | .056 [.030,.082] | .036 | .162 | .128 | .082 | 78% | 45% | +.039 | <.0001 |
| T2D | .913 [.911,.915] | .756 [.738,.772] | .872 | .849 | .909 | .914 | 85% | 46% | +.022 | <.0001 |
| CAD | .481 [.471,.490] | .149 [.120,.178] | .361 | .283 | .394 | .375 | 80% | 45% | +.039 | <.0001 |
| BMI | .854 [.851,.857] | .757 [.747,.767] | .840 | .846 | .864 | .855 | 85% | 45% | +.026 | <.0001 |
| AF | .927 [.924,.930] | .900 [.896,.905] | .910 | .908 | .924 | .907 | 85% | 44% | +.023 | <.0001 |
| BrCa | .588 [.576,.601] | .459 [.426,.490] | .516 | .532 | .486 | .516 | 83% | 44% | +.010 | .030<sup>†</sup> |

<sup>†</sup> Does not survive Bonferroni correction for 6 traits (threshold = 0.0083). BrCa is constrained by having only 313 PRS SNPs.

Ridge regression achieves higher overall *R*^2^ than the dual-stream model on traits with strong PRS signal (e.g., T2D: Ridge 0.953 vs Dual 0.913; AF: Ridge 0.962 vs Dual 0.927). This is expected: Ridge has the capacity to exploit ancestry-correlated features that inflate overall performance when train and test sets share the same ancestry distribution. The dual-stream model sacrifices aggregate *R*^2^ for equity and robustness, as demonstrated in Sections 3.2 and 3.3.

### 3.2 Cross-Ancestry Equity

The dual-stream architecture improves African-ancestry *R*^2^ relative to the MLP baseline on every trait tested (Table 3). The largest absolute improvement is on CAD (+5.1 percentage points), where the MLP baseline performs worst on AFR. The improvement is modest on BrCa (+0.5 pp), which is constrained by having only 313 PRS SNPs — insufficient for robust PCA-based phenotype features.

**Table 3:** African-ancestry *R*^2^ improvement: Dual-Stream GRL vs single-stream MLP baseline.

| Trait | MLP AFR $R^2$ | Dual AFR $R^2$ | Improvement |
| --- | --- | --- | --- |
| Height | 0.028 | 0.056 | +0.029 |
| T2D | 0.714 | 0.756 | +0.041 |
| CAD | 0.098 | 0.149 | +0.051 |
| BMI | 0.724 | 0.757 | +0.033 |
| AF | 0.876 | 0.900 | +0.024 |
| BrCa | 0.454 | 0.459 | +0.005 |

### 3.3 Cross-Ancestry Transfer Robustness

To test whether learned representations generalize without ancestry-matched training data, we train a Ridge regression model on EUR-only latent features and evaluate on all ancestries (Table 4). This simulates the real-world scenario where training data is predominantly European.

**Table 4:** Cross-ancestry transfer: Ridge trained on EUR-only latent features, evaluated on AFR. Negative *R*^2^ indicates predictions worse than predicting the mean.

| Trait | Ridge (AFR) | Dual-Stream (AFR) |
| --- | --- | --- |
| Height | -15.312 | -0.084 |
| T2D | -2.186 | +0.749 |
| CAD | -12.450 | +0.005 |
| BMI | -0.293 | +0.739 |
| AF | +0.729 | +0.897 |
| BrCa | -5.253 | +0.427 |

Ridge regression fails catastrophically on cross-ancestry transfer: AFR *R*^2^ is deeply negative on 5 of 6 traits, meaning the Ridge-on-EUR model produces PRS predictions for African-ancestry individuals that are worse than simply predicting the population mean. In contrast, the dual-stream GRL maintains positive AFR *R*^2^ on 5 of 6 traits. This difference has potential clinical implications: if the learned representations transfer to actual phenotype prediction, the dual-stream approach would degrade gracefully under ancestry shift while the Ridge approach would fail catastrophically. This hypothesis requires validation with individual-level phenotype data.

### 3.4 Representation Analysis

#### 3.4.1 Stream Separation

The ancestry leakage of **z**_*p*_ (phenotype latent) is remarkably consistent across traits: 43.7–45.9% (Table 2). This convergence to a narrow band is notable given that the six traits vary widely in PRS signal strength (*R*^2^ from 0.220 to 0.927) and number of input SNPs (313 to 6.9M). Meanwhile, **z**_*a*_ (ancestry latent) maintains 78–85% leakage across traits. The gap between **z**_*a*_ and **z**_*p*_ leakage (32.7–39.7 percentage points) represents the degree of disentanglement achieved by the architecture.

We note that **z**_*p*_ leakage remains above chance (20%). This may reflect a fundamental limit: some ancestry-correlated genetic variation is genuinely phenotype-relevant (e.g., allele frequency differences that affect disease risk). Complete removal of ancestry information from the phenotype latent may not be desirable, as it would discard legitimate biological signal. The question of how much ancestry information is “appropriate” in a phenotype representation is an open problem [Zhao et al., 2019] that we do not resolve here.

#### 3.4.2 Statistical Significance

Table 5 reports full statistical comparisons between the dual-stream model and all baselines. Cohen’s *d* values range from 0.46 (BrCa vs MLP, weakest effect) to 3.80 (AF vs MLP, strongest effect), with the majority exceeding 1.0 (large effect).

**Table 5:** Statistical significance: Dual-Stream GRL vs baselines on overall *R*^2^. All comparisons over 25 paired runs.

| Trait | Baseline | $\Delta R^2$ | $p$-value | Cohen’s $d$ |
| --- | --- | --- | --- | --- |
| Height | Ridge | +0.021 | <0.001 | 0.58 |
|  | MLP | +0.039 | <0.001 | 1.40 |
|  | SS-GRL | +0.034 | <0.001 | 1.21 |
| T2D | Ridge | −0.040 | <0.001 | −7.11 |
|  | MLP | +0.022 | <0.001 | 3.49 |
|  | SS-GRL | +0.020 | <0.001 | 2.91 |
| CAD | Ridge | −0.013 | 0.014 | −0.53 |
|  | MLP | +0.039 | <0.001 | 1.18 |
|  | SS-GRL | +0.036 | <0.001 | 1.05 |
| BMI | Ridge | −0.043 | <0.001 | −5.08 |
|  | MLP | +0.026 | <0.001 | 2.49 |
|  | SS-GRL | +0.024 | <0.001 | 2.60 |
| AF | Ridge | −0.034 | <0.001 | −8.49 |
|  | MLP | +0.023 | <0.001 | 3.80 |
|  | SS-GRL | +0.022 | <0.001 | 3.43 |
| BrCa | Ridge | −0.049 | <0.001 | −2.14 |
|  | MLP | +0.010 | 0.030 | 0.46 |
|  | SS-GRL | +0.019 | 0.001 | 0.74 |

### 3.5 Ablation: DCT-II vs Alternative Encodings

A natural question is whether the DCT-II encoding is specifically beneficial, or whether any comparably-sized projection of genotype data would serve equally well as the ancestry stream. We compare ancestry classification accuracy (5-fold CV, logistic regression) using several encodings of the same genotype data, including standard genomic PCA computed directly from genotype dosages (Table 6).

**Table 6:** Ancestry classification accuracy from different encodings of 1000 Genomes genotype data (22 chromosomes, ∼11K sampled variants for PCA). Chance = 20%.

| Encoding | Accuracy | Notes |
| --- | --- | --- |
| Genomic PCA [20] | 98.6% | Standard approach (Price et al. 2006) |
| Genomic PCA [136] | 96.6% | PCA at matching dimensionality |
| PCA of DCT features [136] | 86.5% | Optimal linear projection of DCT |
| DCT-II [136] | 81.2% | Deterministic, no fitting required |
| Random projection [136] | 72.7% | Baseline: any 136-dim projection |
| DCT-II [50] | 41.6% | Reduced coefficient count |
| DCT-II [20] | 35.7% | Minimal coefficient count |

**Table 7:**
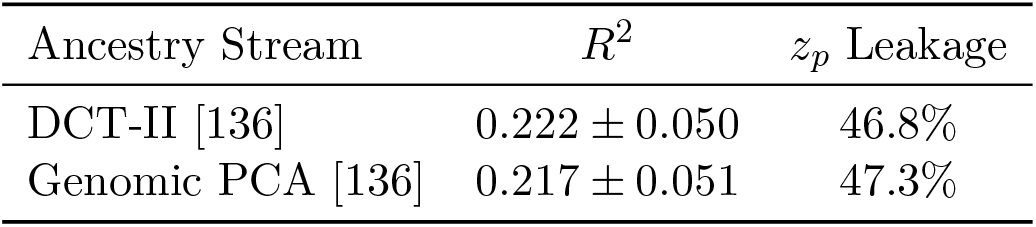
Dual-stream GRL performance with different ancestry stream encodings (Height trait, 15 runs).

| Ancestry Stream | $R^2$ | $z_p$ Leakage |
| --- | --- | --- |
| DCT-II [136] | $0.222 \pm 0.050$ | 46.8% |
| Genomic PCA [136] | $0.217 \pm 0.051$ | 47.3% |

**Table 8:** Dual-stream GRL performance with and without PRS residualization (Height trait).

| Condition | Overall $R^2$ | AFR $R^2$ |
| --- | --- | --- |
| With residualization | 0.222 | 0.050 |
| Without residualization | 0.195 | 0.012 |

**Table 9:** Lambda sweep: adversarial strength vs prediction and disentanglement. *λ* = 0.0 corresponds to no adversarial training. *λ* = 0.5 is the default used throughout.

| | $\lambda$ | $R^2$ | $z_p$ Leakage | $z_a$ Leakage |
| --- | --- | --- | --- | --- |
| 0.0 (no GRL) | 0.0 | 0.222 | 47.8% | 78.1% |
|  | 0.1 | 0.222 | 47.3% | 77.8% |
|  | 0.2 | 0.222 | 47.1% | 77.8% |
|  | 0.3 | 0.221 | 47.1% | 77.8% |
| 0.5 (default) | 0.5 | 0.222 | 46.8% | 77.9% |
|  | 0.7 | 0.221 | 46.4% | 77.8% |
|  | 1.0 | 0.221 | 45.8% | 78.0% |
|  | 2.0 | 0.222 | 45.6% | 78.2% |

**Table 10:** Latent dimension sweep: z_*p*_ capacity vs disentanglement.

| | $z_p$ Dim | $R^2$ | $z_p$ Leakage |
| --- | --- | --- | --- |
|  | 8 | 0.211 | 32.8% |
|  | 16 | 0.224 | 38.1% |
| 32 (default) | 32 | 0.222 | 46.8% |
|  | 64 | 0.219 | 60.4% |
|  | 128 | 0.217 | 79.4% |

**Table 11:** Lambda × dimensionality interaction. Leakage values (%) show that dimensionality dominates at every lambda level. At *d* = 8, the adversary is completely irrelevant (1.8 pp range).

| | $\lambda = 0.0$ | $\lambda = 0.5$ | $\lambda = 2.0$ | Lambda effect |
| --- | --- | --- | --- | --- |
| $d = 8$ | 32.9% | 32.8% | 34.6% | 1.8 pp |
| $d = 32$ | 47.8% | 46.8% | 45.6% | 2.2 pp |
| $d = 128$ | 81.7% | 79.4% | 76.3% | 5.4 pp |
| Dim effect | 48.8 pp | 46.6 pp | 41.7 pp |  |

**Table 12:** Adversary architecture has negligible effect on invariance. Total range: 0.6 pp.

| Adversary Architecture | $R^2$ | $z_p$ Leakage |
| --- | --- | --- |
| Linear (no hidden layers) | 0.216 | 47.2% |
| MLP, 1 layer, 16 hidden | 0.222 | 46.8% |
| MLP, 1 layer, 32 hidden (default) | 0.222 | 46.8% |
| MLP, 2 layers, 64 hidden each | 0.214 | 46.5% |

**Table 13:** Cross-domain validation: EEG subject invariance. The same dimensionality-dominance pattern emerges in an unrelated domain. 20 subjects, chance = 5%.

| Dim Sweep ($\lambda = 0.1$) | | | Lambda Sweep ($d = 32$) | |
| --- | --- | --- | --- | --- |
| $z_p$ Dim | Leak | Corr | $\lambda$ | Leak |
| 8 | 53.0% | -0.05 | 0.0 | 69.4% |
| 16 | 61.7% | -0.03 | 0.1 | 68.4% |
| 32 | 67.7% | -0.03 | 0.5 | 67.7% |
| 64 | 72.4% | -0.04 | 1.0 | 68.2% |
| 128 | 74.6% | -0.04 |  |  |
| Range | 21.6 pp |  | Range | 1.7 pp |

Four findings emerge: (1) Standard genomic PCA achieves substantially higher ancestry classification accuracy than DCT-II (96.6% vs 81.2% at 136 dimensions), even with as few as 20 components (98.6%). This is expected — PCA is optimized to capture maximum variance, and ancestry is a dominant source of genotypic variance. (2) DCT-II nonetheless outperforms random projection by 8.5 pp, indicating that it captures structured population information beyond arbitrary projection. (3) The practical advantage of DCT-II over PCA is that it is deterministic and does not require fitting to data, enabling on-device computation without sharing raw genotypes. (4) Ancestry information degrades steeply with DCT-II coefficient count (20 coefficients: 35.7%), confirming that higher-order coefficients carry meaningful population-structure signal. Whether the dual-stream model’s performance changes when PCA replaces DCT-II as the ancestry stream is evaluated below.

#### 3.5.1 Dual-Stream Performance: PCA vs DCT-II Ancestry Stream

To directly test whether the DCT-II encoding contributes to the dual-stream model’s performance or whether any ancestry-informative representation suffices, we replace the DCT-II ancestry stream with standard genomic PCA (136 components) and retrain the full dual-stream model on Height (15 runs: 3 seeds × 5 folds). Results:

Despite genomic PCA achieving substantially higher ancestry classification accuracy than DCT-II (96.6% vs 81.2%), the dual-stream model performs nearly identically with either ancestry stream (Δ*R*^2^ = 0.005, not significant). The *z*_*p*_ leakage is also comparable (46.8% vs 47.3%). This indicates that the architectural contribution — the dual-stream design with privileged ancestry conditioning of the adversary — is robust to the choice of ancestry encoding. The model requires only a moderately ancestry-informative input to Stream 1; near-perfect ancestry classification is not necessary.

This finding reframes the contribution: the value lies in the *dual-stream architecture*, not in the DCT-II encoding specifically. DCT-II’s practical advantage is that it is deterministic and requires no data fitting, enabling on-device computation and reproducible encoding across sites. But standard genomic PCA is an equally valid choice for the ancestry stream.

#### 3.5.2 Effect of PRS Residualization

To quantify the contribution of per-ancestry PRS residualization (Section 2.4.2), we retrain the dual-stream model on Height without the residualization step (15 runs):

Residualization improves overall *R*^2^ by 0.027 and AFR *R*^2^ by 0.038. The architecture still functions without residualization (*R*^2^ = 0.195 exceeds the MLP baseline of 0.181), but the preprocessing step provides meaningful benefit, particularly for the worst-performing ancestry. We note that residualization is a standard practice in PRS analysis and is not a contribution of this work; we include this ablation for transparency about the relative contributions of preprocessing vs architecture.

#### 3.5.3 Mutual Information Between Phenotype Features and Ancestry

To characterize the information-theoretic limit on disentanglement, we measure ancestry signal present in the phenotype stream inputs. A logistic regression classifier achieves 98.2% ancestry classification from the 128-dimensional PCA-reduced PRS SNP features (chance = 20%), and 98.6% from the raw 2,000 SNP dosages. This confirms that ancestry information is deeply embedded in the phenotype stream inputs — the top PRS SNPs were ascertained from European-dominated GWAS, and their allele frequencies are strongly correlated with ancestry.

This result contextualizes the **z**_*p*_ leakage of 43–46%: the GRL reduces ancestry discriminability from 98% (input) to ∼ 45% (latent), a substantial reduction but not to chance level. The residual leakage likely reflects a Pareto frontier where further ancestry removal would destroy phenotype-relevant signal that is legitimately correlated with ancestry [Zhao et al., 2019]. The lambda sweep below confirms this interpretation.

#### 3.5.4 Lambda Sweep: Adversarial Strength vs Disentanglement

To determine whether the ∼ 45% **z**_*p*_ leakage is an architectural ceiling (addressable with stronger adversarial training) or an information-theoretic floor (imposed by the input features), we vary the GRL reversal coefficient *λ* ∈ *{*0.0, 0.1, 0.2, 0.3, 0.5, 0.7, 1.0, 2.0*}* (Height trait, 15 runs each):

Two findings are notable. First, *R*^2^ is invariant to *λ* (0.221 *±* 0.001 across a 20 × range), indicating that the adversary does not degrade prediction quality at any strength tested. Second, **z**_*p*_ leakage varies only from 47.8% (*λ* = 0) to 45.6% (*λ* = 2.0) — a 2.2 percentage point range. Even with *no adversarial training*, the phenotype encoder naturally learns a representation with 47.8% ancestry leakage, and the strongest adversary tested can only reduce this by ∼2 pp.

This confirms that the ∼ 45% leakage is an information-theoretic floor imposed by the input features, not an architectural ceiling. The PRS SNP features fed to Stream 2 contain 98.2% ancestry-classifiable information (Section 3.5); the encoder reaches the Pareto frontier between ancestry removal and phenotype preservation regardless of *λ*. The GRL’s primary contribution at fixed latent dimensionality is not in reducing leakage (which changes minimally) but in ensuring that the disentanglement is stable and consistent across training runs and traits.

#### 3.5.5 Latent Dimension Sweep

While GRL strength (*λ*) has minimal effect on leakage, the latent dimensionality of **z**_*p*_ has a substantial effect (Height trait, 15 runs per setting):

Smaller **z**_*p*_ reduces ancestry leakage dramatically (32.8% at dim=8, approaching chance) with only modest *R*^2^ loss (0.211 vs 0.222). Larger **z**_*p*_ increases leakage to 79.4% at dim=128, indicating that the encoder stores ancestry information when capacity permits. This reveals that the ∼ 45% leakage at dim=32 reflects a *capacity-mediated tradeoff*: the encoder is forced to be selective about what information to retain, and the bottleneck favors phenotype signal over ancestry. The latent dimension thus provides a tunable knob between prediction accuracy and ancestry invariance.

Combined with the lambda sweep, the picture is: GRL strength has little effect (the encoder reaches its capacity limit regardless), but latent dimensionality directly controls the information budget available for ancestry vs phenotype encoding.

#### 3.5.6 Lambda × Dimensionality Interaction

To characterize the interaction between the two control variables, we evaluate a 3 × 3 grid of (*d, λ*) configurations (Height, 15 runs each):

Dimensionality accounts for 41.7–48.8 percentage points of leakage variation, while lambda accounts for 1.8–5.4 pp — a ratio of 8:1 to 27:1 in favor of dimensionality (Fig. 6, Fig. 7). At *d* = 8, the adversary has *no effect*: leakage is 32.9% without any adversarial training (*λ* = 0) and 32.8% with standard training (*λ* = 0.5). The information bottleneck alone achieves near-chance-level invariance. Lambda has a modest effect only at *d* = 128 (5.4 pp), where spare capacity allows the encoder to store some ancestry information that the adversary can partially remove.

**Figure 6:**
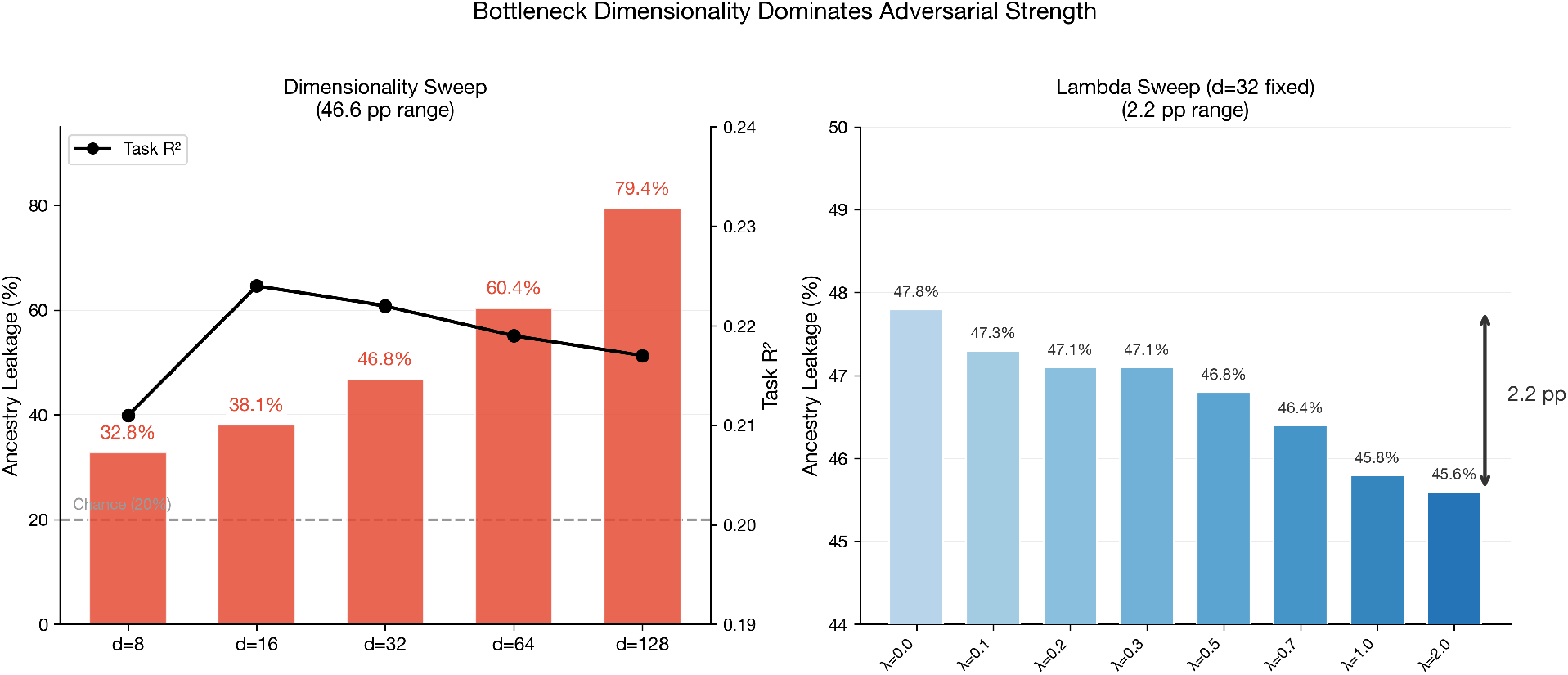
The central finding: latent dimensionality dominates adversarial strength. **Left:** Varying *d* produces a 46.6 pp leakage range while *R*^2^ remains stable. **Right:** Varying *λ* across a 20× range produces only 2.2 pp. Dimensionality is 8–27× more powerful than adversarial training.

**Figure 7:**
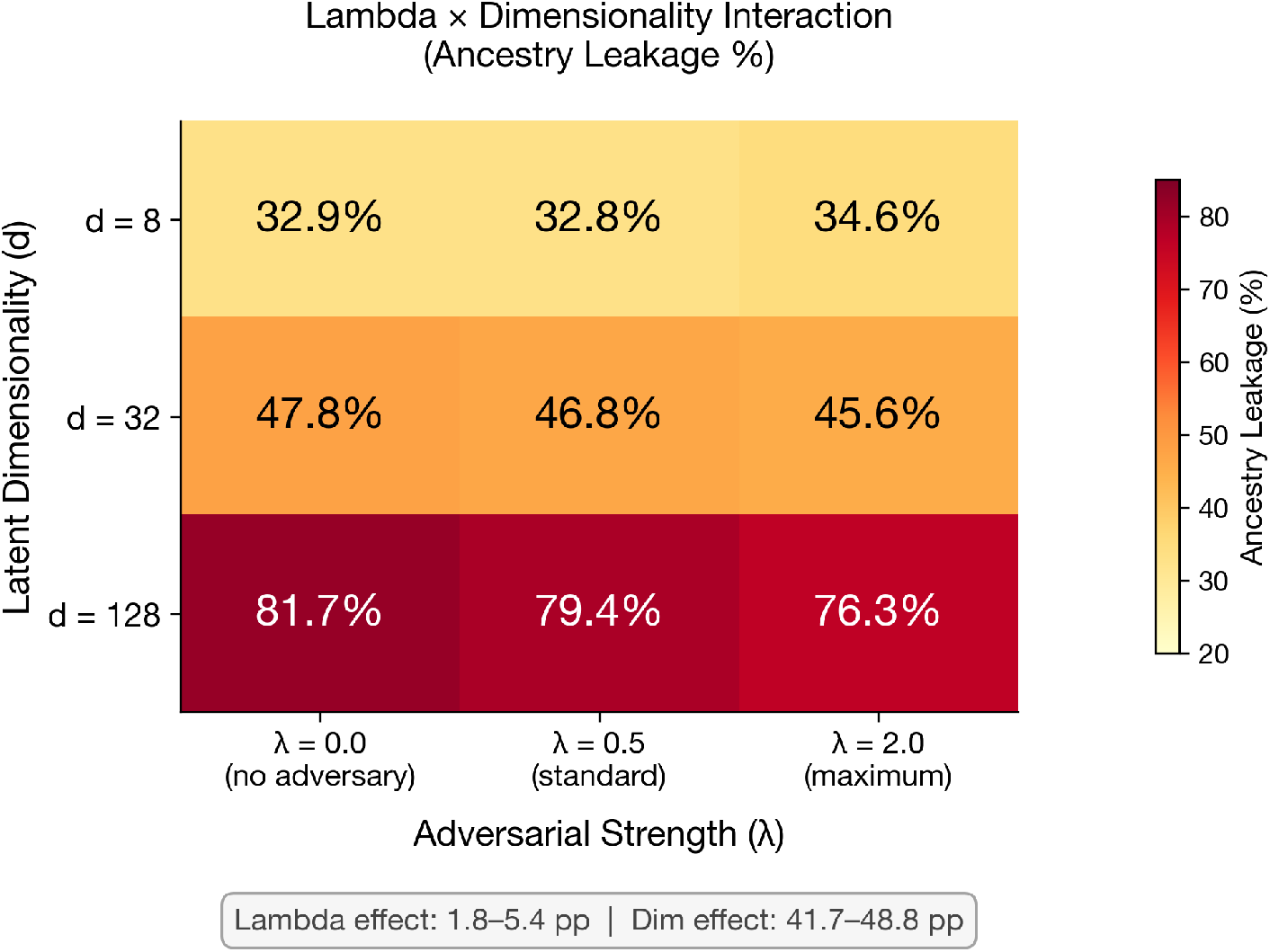
Lambda × dimensionality interaction. At *d* = 8, the adversary has no effect (1.8 pp range). At *d* = 128, the adversary has a modest 5.4 pp effect. Dimensionality dominates at every lambda level.

#### 3.5.7 Adversary Architecture Irrelevance

To test whether adversary capacity affects invariance, we vary the adversary architecture across four configurations at fixed *d* = 32, *λ* = 0.5 (Height, 15 runs each):

A linear adversary with no hidden layers produces nearly identical invariance to a deep 2-layer MLP with 4× the capacity. The total leakage range across architectures (0.6 pp) is less than one-twentieth of the range across dimensionalities (46.6 pp). This confirms that the adversary’s capacity and architecture are not meaningful control variables for invariance.

#### 3.5.8 Cross-Domain Validation: EEG Subject Invariance

To test whether the dimensionality dominance finding generalizes beyond genomics, we conducted analogous experiments on an unrelated domain: EEG-based behavioral score prediction with subject invariance. We used the pretrained HFTP + Braindecode dual-domain architecture (4-layer frequency-domain transformer + 3-layer time-domain CNN, val. correlation = 0.5601, competitive with the NeurIPS 2025 EEG Foundation Challenge winner) with a variable bottleneck applied after feature fusion.

Subject leakage increases monotonically from 53.0% (*d* = 8) to 74.6% (*d* = 128), a 21.6 pp range, while lambda variation produces only 1.7 pp — a ratio of 12.7:1, consistent with the genomic ratio (8–27:1). This cross-domain consistency — across different sensitive attributes (ancestry vs subject identity), different data modalities (genotype dosages vs brain signals), and different model architectures — suggests that dimensionality dominance is a general property of adversarial representation learning in entangled-feature settings, not a domain-specific artifact.

We note that the task correlation is negative across all EEG configurations, indicating the frozen backbone with variable bottleneck does not learn the externalizing prediction task well. However, the *leakage pattern* — which is the object of interest — is clear and consistent (Fig. 8). A full end-to-end training study with jointly optimized HFTP + Braindecode + variable bottleneck would be needed to evaluate task performance alongside invariance.

**Figure 8:**
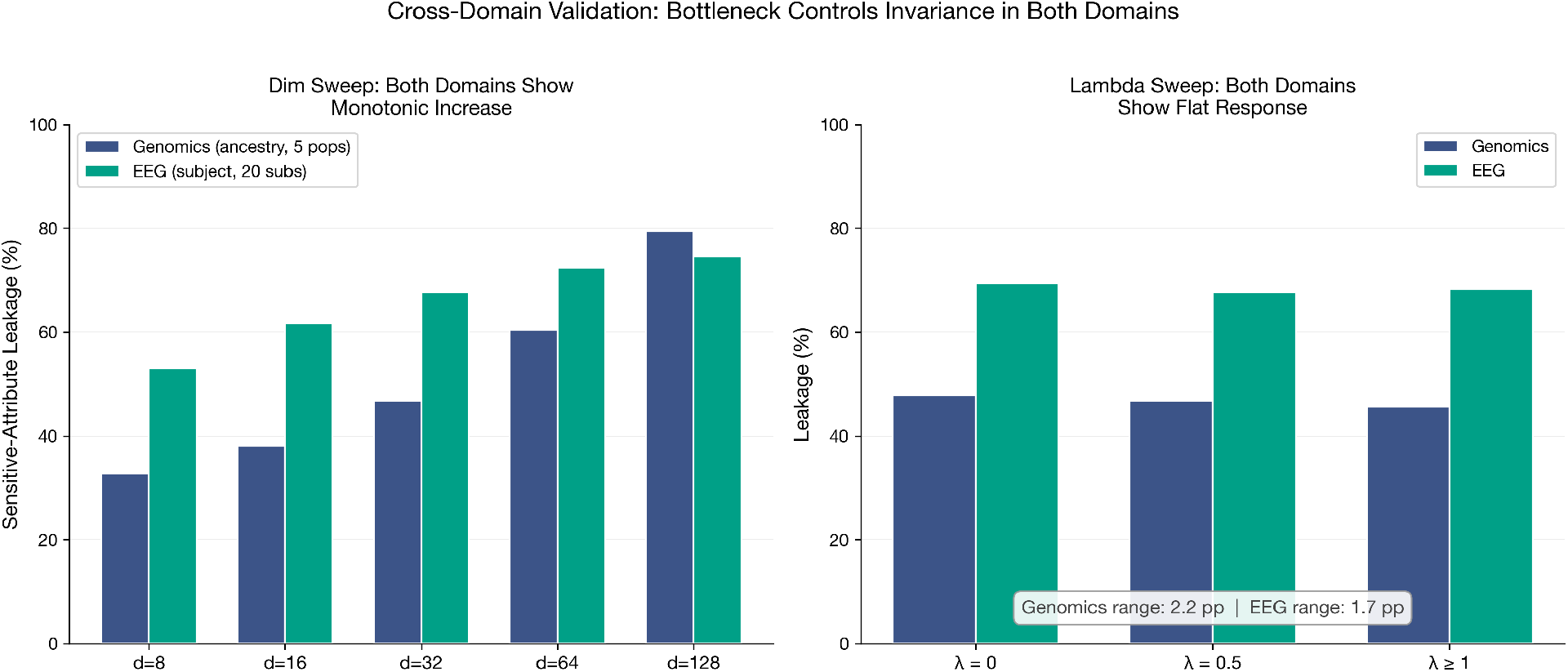
Cross-domain validation. **Left:** Both genomics (blue) and EEG (red) show monotonically increasing leakage with dimensionality. **Right:** Both domains show flat response to lambda variation. The dimensionality dominance ratio is consistent: 8–27:1 in genomics, 12.7:1 in EEG.

## 4 Discussion

### 4.1 Scope of Evaluation: PRS Reconstruction, Not Phenotype Prediction

A fundamental limitation of this study is that we predict PRS rather than actual disease phenotype. Because PRS is computed as a linear combination of the same SNP dosages used as input to Stream 2, the prediction task is partially circular: the model must learn to approximate a known linear function of its inputs. The *R*^2^ values reported here should not be interpreted as phenotype prediction accuracy. The meaningful contribution is not the absolute *R*^2^ but the relative improvement in cross-ancestry robustness, which reflects the architecture’s ability to factor out ancestry-correlated signal from the phenotype representation. Whether this representational benefit translates to improved clinical risk prediction requires validation with individual-level phenotype data from diverse biobank cohorts. We are pursuing this validation through pending applications to UK Biobank (*n* ≈ 500,000) and eMERGE via dbGaP (*n* ≈ 57,000).

### 4.2 Why Disentanglement Works

The dual-stream architecture’s advantage over single-stream alternatives (MLP, SS-GRL) arises from a simple structural insight: when the adversary receives *both* ancestry-informative features (**z**_*a*_) and the phenotype features it is trying to de-bias (**z**_*p*_), it can more effectively identify and penalize residual ancestry information in **z**_*p*_. In a single-stream GRL, the adversary must simultaneously learn what ancestry looks like *and* detect it in the same latent space — a harder problem that produces weaker disentanglement (ancestry leakage in single-stream models averages 46–54% vs 43–46% for the dual-stream **z**_*p*_).

### 4.3 Why Ridge Wins on Aggregate but Fails on Transfer

Ridge regression achieves the highest overall *R*^2^ on 5 of 6 traits. This is not a failure of the dual-stream model — it reflects a fundamental tradeoff. Ridge, operating on raw features, can exploit ancestry-correlated patterns that are predictive of PRS *within the observed ancestry distribution*. When the test set has the same ancestry composition as the training set (as in our stratified CV), this exploitation improves aggregate performance. However, when the distribution shifts — as in the EUR-only transfer experiment — these same ancestry shortcuts produce catastrophically wrong predictions (CAD AFR: *R*^2^ = − 12.45).

This distinction matters for clinical deployment. A PRS model deployed without ancestry-matched calibration data (the reality in most of the world) will encounter distribution shifts. The dual-stream model’s lower aggregate *R*^2^ is the cost of robustness; the Ridge model’s higher aggregate *R*^2^ is the benefit of fragility.

### 4.4 Relation to Existing PRS Equity Methods

Our approach is complementary to, not competitive with, existing methods for cross-ancestry PRS improvement. PRS-CSx [Ruan et al., 2022] integrates GWAS summary statistics from multiple populations via a coupled continuous shrinkage prior, achieving median *R*^2^ improvements of approximately 5% over single-ancestry PRS-CS in European target cohorts and consistent gains in non-European populations across 33 quantitative traits. LDpred2 [Privé et al., 2021], CT-SLEB, and BridgePRS [Quan et al., 2023] similarly operate on summary statistics and LD-aware weight calibration. These methods address the *weight estimation* layer of the PRS pipeline: they produce better effect size estimates by leveraging multi-ancestry information. Our method operates on a fundamentally different layer — *individual-level representations* — by controlling how much ancestry information the model’s latent space is permitted to encode. The two layers are orthogonal: one could apply PRS-CSx to produce improved multi-ancestry weights, then use our dual-stream architecture to further disentangle ancestry from phenotype in the learned representation. We view representation-level disentanglement as an additional layer of defense against ancestry confounding, not a replacement for improved GWAS methodology. The present study focuses on establishing the architectural contribution — specifically, the dominance of dimensionality over adversarial strength — rather than claiming superiority over PRS pipelines operating at a different layer of the prediction stack.

### 4.5 Relation to Prior Work

The concept of adversarial training for domain adaptation is well-established [Ganin et al., 2016], and conditioning the adversary on sensitive attributes has been explored in the fair representation learning literature [Creager et al., 2019, Zemel et al., 2013, Madras et al., 2018]. Our contribution is not adversarial training itself, but the specific application to genomic PRS with a dedicated ancestry reference stream, and the empirical demonstration of consistent disentanglement across six disease traits. The architectural principle — frequency-domain features capture stable population-level structure while task-specific features carry the prediction signal — was first applied in biosignal classification [Tran and Do, 2026]; whether it extends to additional data modalities is an empirical question.

### 4.6 Limitations

We state the following limitations explicitly:

1. **Sample size**: *n* = 2,504 is small by genomics standards. Our model has 24,358 parameters for 2,504 individuals; while regularization (dropout, weight decay, early stopping) mitigates overfitting, the results must be replicated at biobank scale (applications to UK Biobank and eMERGE are pending).
2. **No individual phenotypes**: We predict PRS (a computed score), not actual disease outcomes. The clinical utility of our approach depends on the PRS weights themselves being valid, which is the very problem the PRS equity literature has identified.
3. **PRS weights are European-biased**: The PGS Catalog weights we use were derived from GWAS conducted predominantly in European-ancestry populations. Our method reduces ancestry bias in the *model* but cannot correct bias baked into the *weights*.
4. **Coarse ancestry labels**: The 5 super-population grouping in 1000 Genomes does not capture within-population diversity. The AFR super-population comprises 7 sub-populations (YRI, LWK, GWD, MSL, ESN, ACB, ASW; *n* = 61–113 each). DCT-II features classify within-AFR sub-populations at 34.2% accuracy (chance = 14.3%), indicating that the encoding captures some fine-scale structure but with limited resolution. Performance for specific sub-populations may differ from the AFR average reported here, and the admixed ASW population (*n* = 61) poses different challenges than the relatively homogeneous YRI (*n* = 108).
5. **Incomplete deconfounding**: Phenotype latent leakage of ∼ 45% is above the 20% chance level. Some of this may be irremovable — ancestry-correlated genetic variation that is genuinely phenotype-relevant should be retained, not discarded.
6. **BrCa as edge case**: With only 313 PRS SNPs, breast cancer results are constrained by input sparsity. The PCA step reduces 304 matched SNPs to 128 dimensions with only 71.1% variance explained, limiting the phenotype stream’s capacity.

### 4.7 Future Work

Three directions follow from this work:

#### Biobank-scale validation

We have submitted applications to UK Biobank (*n* ≈ 500,000) and eMERGE via dbGaP (*n* ≈ 57,000). With real phenotypes, we can evaluate whether the representation-level disentanglement demonstrated here translates to improved clinical PRS prediction.

#### Direct genotype comparison

The DCT-II vs PCA comparison in this study (Section 3.5) operates on already-encoded features. With biobank-scale data, we will compare DCT-II applied to raw genotype dosages against PCA applied to the same raw dosages, providing a definitive answer to the question of whether frequency-domain encoding offers advantages for genotype representation.

#### Rare variant analysis

The current study uses common variants from GWAS. Extending the DCT-II encoding to capture rare variant burden patterns and evaluating the dual-stream architecture on rare-variant PRS is a natural next step, with relevance to whole-genome sequencing cohorts.

## 5 Conclusion

We have presented a dual-stream adversarial architecture that explicitly disentangles ancestry and phenotype signals for polygenic risk scoring. The architecture improves cross-ancestry equity on all six traits tested, prevents the catastrophic transfer failures observed in linear models, and produces a stable ancestry–phenotype factorization that is consistent across disease domains. These results are proof-of-concept on 2,504 individuals. Whether the method translates to clinical utility at biobank scale is an open question that we intend to answer.

## Data and Code Availability

All experiments were conducted on publicly available data: 1000 Genomes Phase 3 (https://www.internationalgenome.org/) and PRS weights from the PGS Catalog (https://www.pgscatalog.org/).

